# High-throughput identification of RNA nuclear enrichment sequences

**DOI:** 10.1101/189654

**Authors:** Chinmay J Shukla, Alexandra L McCorkindale, Chiara Gerhardinger, Keegan D Korthauer, Moran N Cabili, David M Shechner, Rafael A Irizarry, Philipp G Maass, John L Rinn

**Affiliations:** Department of Stem Cell and Regenerative Biology, Harvard University, Cambridge, Massachusetts 02138, USA; Broad Institute of MIT and Harvard, Cambridge, Massachusetts 02139, USA; Department of Biostatistics and Computational Biology, Dana-Farber Cancer Institute, Cambridge, Massachusetts 02115, USA; Program in Biological and Biomedical Sciences, Harvard Medical School, Boston, Massachusetts 02115, USA; Department of Biostatistics, Harvard T.H. Chan School of Public Health, Boston, Massachusetts 02115, USA; Department of Pathology, Beth Israel Deaconess Medical Center, Boston, Massachusetts 02215, USA; Berlin Institute for Medical Systems Biology, Max Delbrück Center for Molecular Medicine, Berlin-Buch 13125, Germany

## Abstract

One of the biggest surprises since the sequencing of the human genome has been the discovery of thousands of long noncoding RNAs (lncRNAs)^1–6^. Although lncRNAs and mRNAs are similar in many ways, they differ with lncRNAs being more nuclear-enriched and in several cases exclusively nuclear^7,8^. Yet, the RNA-based sequences that determine nuclear localization remain poorly understood^9–11^. Towards the goal of systematically dissecting the lncRNA sequences that impart nuclear localization, we developed a massively parallel reporter assay (MPRA). Unlike previous MPRAs^12–15^ that determine motifs important for transcriptional regulation, we have modified this approach to identify sequences sufficient for RNA nuclear enrichment for 38 human lncRNAs. Using this approach, we identified 109 unique, conserved nuclear enrichment regions, originating from 29 distinct lncRNAs. We also discovered two shorter motifs within our nuclear enrichment regions. We further validated the sufficiency of several regions to impart nuclear localization by single molecule RNA fluorescence *in situ* hybridization (smRNA-FISH). Taken together, these results provide a first systematic insight into the sequence elements responsible for the nuclear enrichment of lncRNA molecules.

---

RNA subcellular localization provides a fundamental mechanism through which cells modulate and utilize the functions encoded in their transcriptomes^16^. This spatial layer of gene regulation is known to be critical in a variety of contexts, including asymmetric cell divisions^17^, embryonic development^18–20^, and signal transduction^21^. Previous work has identified a small number of *cis*-acting mRNA localization elements, using genetic approaches or hybrid reporter constructs to decipher sequences required for localization to different parts of the cell^16,18^. These elements are often located in 3′ untranslated regions (UTRs), and range from five to several hundred nucleotides in length^9–11,18^. Yet, the sequences and structures responsible for RNA localization remain inchoate. In contrast to mRNAs that are mostly localized outside the nucleus, lncRNAs are enriched or retained in the nucleus. Increasing evidence suggests that many lncRNAs may reside in the nucleus for the purpose of regulating nuclear processes, including formation of paraspeckles, topological organization of the nucleus, and regulation of gene expression^1,3,4,22^. However, while it is now evident that lncRNAs have important functions in the nucleus^22^, very little is known about specific sequence elements driving their nuclear enrichment^9–11^.

To elucidate which sequences drive lncRNA nuclear enrichment, we developed a high-throughput approach for identifying nuclear enrichment elements. Our approach, derived from a massively parallel reporter assay (MPRA)^12–15, 23^ is based on a previous assay demonstrating that the native cytosolic localization of a noncoding RNA reporter (a frame-shifted *Sox2* mutant, “fsSox2”) can be altered by appending this reporter with additional RNA sequences^9^. The MPRA we designed highly parallelizes this assay by appending thousands of oligos to fsSox2. Briefly, we selected 38 lncRNAs with diverse subcellular localization patterns: from single nuclear foci (*e.g. XIST, ANRIL, ANCR, PVT1, KCNQ1OT1, FIRRE*) to broadly diffuse cytosolic patterns (*e.g*. NR_024412, XL0C_012599)^24^. We generated a pool of 11,969 oligos 153 nucleotides in length, each with a unique barcode, that tiles each of the 38 lncRNAs. This pool was expressed in HeLa cells followed by nuclear isolation and targeted deep sequencing to determine the partitioning of each fsSox2 variant (Figure 1A, Extended Data Table 1, *Methods*). All experiments were performed as six-biological replicates to ensure sufficient statistical power for our analytical model, and accurately estimate in-group variance (*see below*, *Methods*).

**Figure 1.**
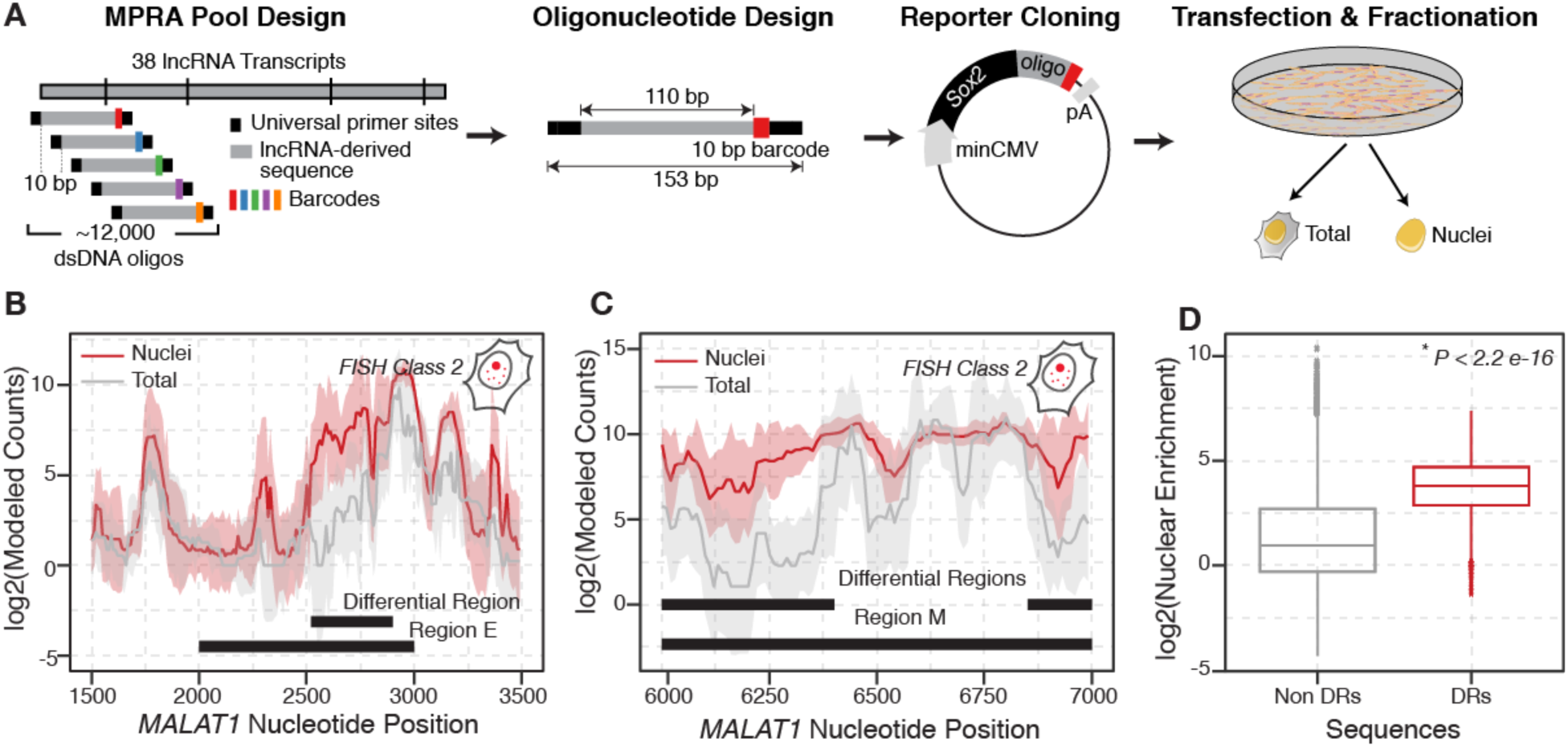
A Massively Parallel Reporter Assay to identify RNA nuclear enrichment signals. **A.** Experimental overview. *Far left:* oligonucleotide pool design. Double-stranded DNA (dsDNA) oligonucleotides were designed by computationally scanning 38 parental lncRNA transcripts (“lncRNA cDNAs,” Extended Datat Table 1) in 110 bp windows, with 10 bp spacing between sequential oligos. These lncRNA-derived “Variable Sequences” (gray) were appended with unique barcodes and primer binding sides, resulting in a pool of 11,969 oligos (**Supplementary Data 1**). The vertical lines in the lncRNA denote splice junctions. *Second from left:* schematic summarizing the design of each pool oligonucleotide. *Second from right:* Reporter design. The oligonucleotide pool was cloned into a reporter plasmid as a transcriptional fusion at the 3'- terminus of the fsSox2 gene. pA: polyadenylation sequence *Far right:* MPRA workflow. The Sox2~oligo reporter pool is transiently transfected into HeLa cells. Following 48h of expression, cells are subsequently fractionated to isolate nuclei, and the nuclear enrichment of each pool member is quantified by targeted RNA sequencing (*not shown*). Matched whole-cell lysates from unfractionated cells serve as controls. **B-C**. Differential Region-calling correctly identifies nuclear retention elements in *MALAT1*. Solid lines: per-nucleotide abundances in the nuclear (red) and whole-cell (gray) fractions, modeled for each position along the *MALAT1* transcript, based on the aggregate behavior of all oligos containing that nucleotide (*Methods*). Shaded regions: standard deviations. Median values for six biological replicates are shown. **D.** Boxplot comparing the nuclear enrichment for all nucleotides within differential regions (“DRs”), relative to all the other nucleotides surveyed (“Non DRs”). *P*-value: Mann Whitney Test.

To identify lncRNA nuclear enriched regions we implemented a statistical method that merges individual nucleotides into longer aggregate regions^25^. We further ranked candidate regions using a newly defined summary statistic, that generates a null distribution for this statistics by permuting sample labels, and uses this null distribution to assigns *p*-values (Extended Data Figure 1; *Methods*). Our approach leverages the inter-replicate variability inherent in high throughput reporter assays and allows us to sensitively and accurately discover nuclear enriched RNA segments which we term “differential regions” (DRs). Importantly, our method allows us to identify DRs greater than individual oligos based on their coherence across larger regions.

To test the performance of our assay and analytic method we first focused on a well established nuclear lncRNA *MALAT1*. Previous work demonstrated that two elements termed Region E and Region M, derived from the lncRNA *MALAT1*, are particularly potent RNA nuclear localization signals^11^. We examined the nuclear enrichment of all fsSox2 pool variants bearing elements derived from lncRNA *MALAT1* (*Methods*). Consistent with the previous study, nucleotides derived from Region E and Region M were highly enriched in the nucleus compared to those residing elsewhere in the human *MALAT1* lncRNA. Thus, our assay can recapitulate known RNA localization signals and our analysis approach can identify localization domains longer than a given tiled oligos.

Next we sough to agnostically and systematically investigate nuclear enrichment regions harbored within 38 lncRNAs. Our analysis identified 109 DRs (FDR < 0.1) originating from 29 distinct lncRNAs that were significantly enriched in nuclear fractions, relative to whole cell lysates (Extended Data Table 2). Two of these DRs overlap and subsume Region M while another DR overlaps with Region E within the *MALAT1* lncRNA (Figure 1B, 1C). To confirm that our approach was robust, we compared the significant DRs to all other regions represented in our pool and found them significantly more nuclear enriched (Figure 1D; *P* < 1/10^6^, Mann-Whitney Test; *Methods*). The localization patterns of the selected 38 lncRNAs have been previously parsed into five smRNA-FISH classes^24^. These included lncRNAs strictly nuclear (FISH Class I), those that are diffusely localized in the cytoplasm (FISH Class V), and three intermediate classes (FISH Classes II-IV). Our MPRA approach discovered DRs derived from lncRNAs in all five FISH classes (Figure 2A–E). Notably, the number of DRs within each class broadly correlated with the degree of nuclear localization observed by smRNA-FISH (Figure 2F). Many strictly-nuclear lncRNAs (FISH Class I) harbor multiple DRs, possibly indicating the presence of a redundant nuclear localization motif. For example, we discovered 18 DRs in *XIST* and 10 DRs in *MALAT1* and some of the DRs we discovered in *XIST* overlap with previously-described repeat elements – RepC and RepD.

**Figure 2.**
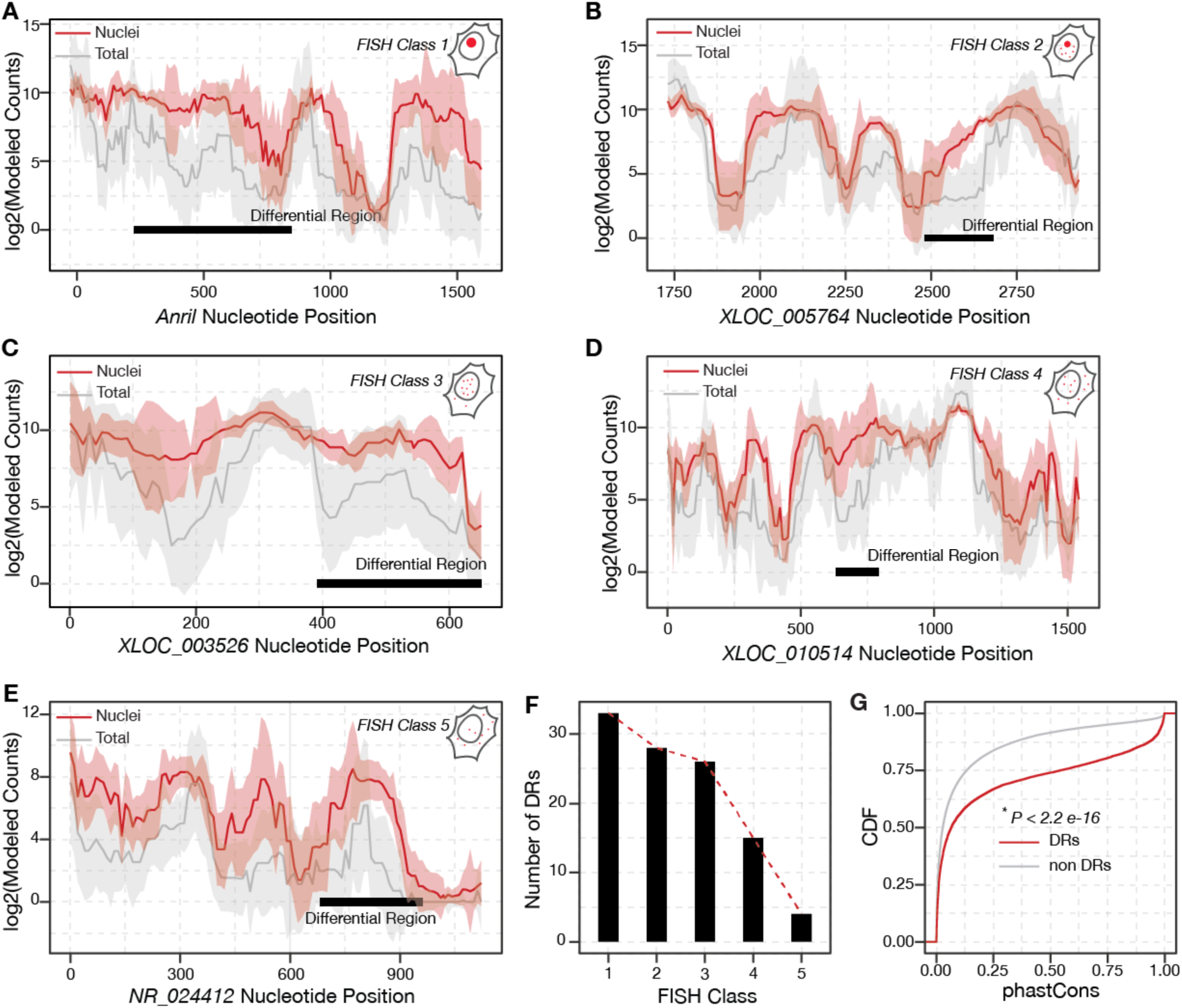
Novel lncRNA nuclear enrichment signals. **A-E**. Identification of Differential Regions within lncRNAs with different subcellular localization patterns. Data are depicted as in Figure 1C. Established subcellular localization patterns range from: **A**. those occupying a single, prominent nuclear focus (*ANRIL*, FISH Class 1), to: **E**. those exhibiting a diffuse, mostly cytosolic pattern (*NR*_024412, FISH Class 5)^24^. **F**. The number of Differential Regions discovered within lncRNAs from each FISH Class correlates with that class’s degree of nuclear localization. **G.** Differential Regions are more highly conserved than are most lncRNA sequences. Cumulative distribution function (CDF) of phastCons scores comparing nucleotides within Differential Regions (*red*), to all other nucleotides within the oligo pool (*gray*). *P*-value: Mann Whitney Test.

We further analyzed the evolutionary conservation, length distribution, and sequence content of these putative nuclear localization sequences. We used phastCons^26, 27^ scores to assess evolutionary conservation, and we observed significantly higher scores among our DRs than in other lncRNA regions tiled by our MPRA (Figure 2G; *P* < 1/10^6^, Mann-Whitney Test; *Methods*). The lengths of our DRs ranged from 80–740 nucleotides (nt), with an average of 300 nt (Extended Data Figure 6A). While we detected a weak correlation between the length of a given lncRNA and number of DRs within (Extended Data Figure 6B), this analysis is confounded by inconsistent length of lncRNAs across the five FISH classes. Finally, we did not observe a difference in GC content between the DRs and other sequences in our tiled lncRNAs (Extended Data Figure 6C).

We hypothesized that our DRs might harbor common sequence motifs or protein-binding preferences. To test this, we searched for motifs that were more prevalent among the DRs than in other regions of the lncRNAs, using the MEME software package^28^. We identified a 57 nt motif occuring 18 times exclusively in *XIST*, and not elsewhere in the human genome (Figure 3A-C). Another, 15 nt “C-rich” motif was found in 52 DRs of 21 different lncRNAs (Figure 3D–F), and we discovered four additional motifs closely related to the described here (Extended Data Figure 7A–D). Similarly, k-mer analysis^29^ revealed several C-rich 4-mers that were mildly predictive of a DR (Extended Data Figure 7E). In total, we discovered six motifs and confirmed that the nucleotides overlapping these motifs were significantly enriched in the nucleus (*P* < 1/10^6^, Mann-Whitney Test, *Methods)*, compared to all other regions tiled in our MPRA (Figure 3G). Since the C-rich motif occured in more than 50 distinct DRs of diverse lncRNAs, we postulated that this motif could function as a global RNA nuclear localization element. To test this, we examined the nuclear-cytoplasmic localizations of all human transcripts containing this motif, using fractionation RNA-Seq data from ENCODE^30^. We observed a modest increase (*P* < 1/10^6^, Mann-Whitney Test) in nuclear localization of transcripts with the C-rich motifs across all 11 ENCODE TIER 2 cell lines (Figure 3H, I, Extended Data Figure 8). This further demonstrates the potential power of our MPRA to discover functional elements that may be missed by classic RNA localization studies. A similar C-rich motif was recently discovered by another group and has been investigated in mechanistic detail (Igor Ulitsky – personal communication).

**Figure 3.**
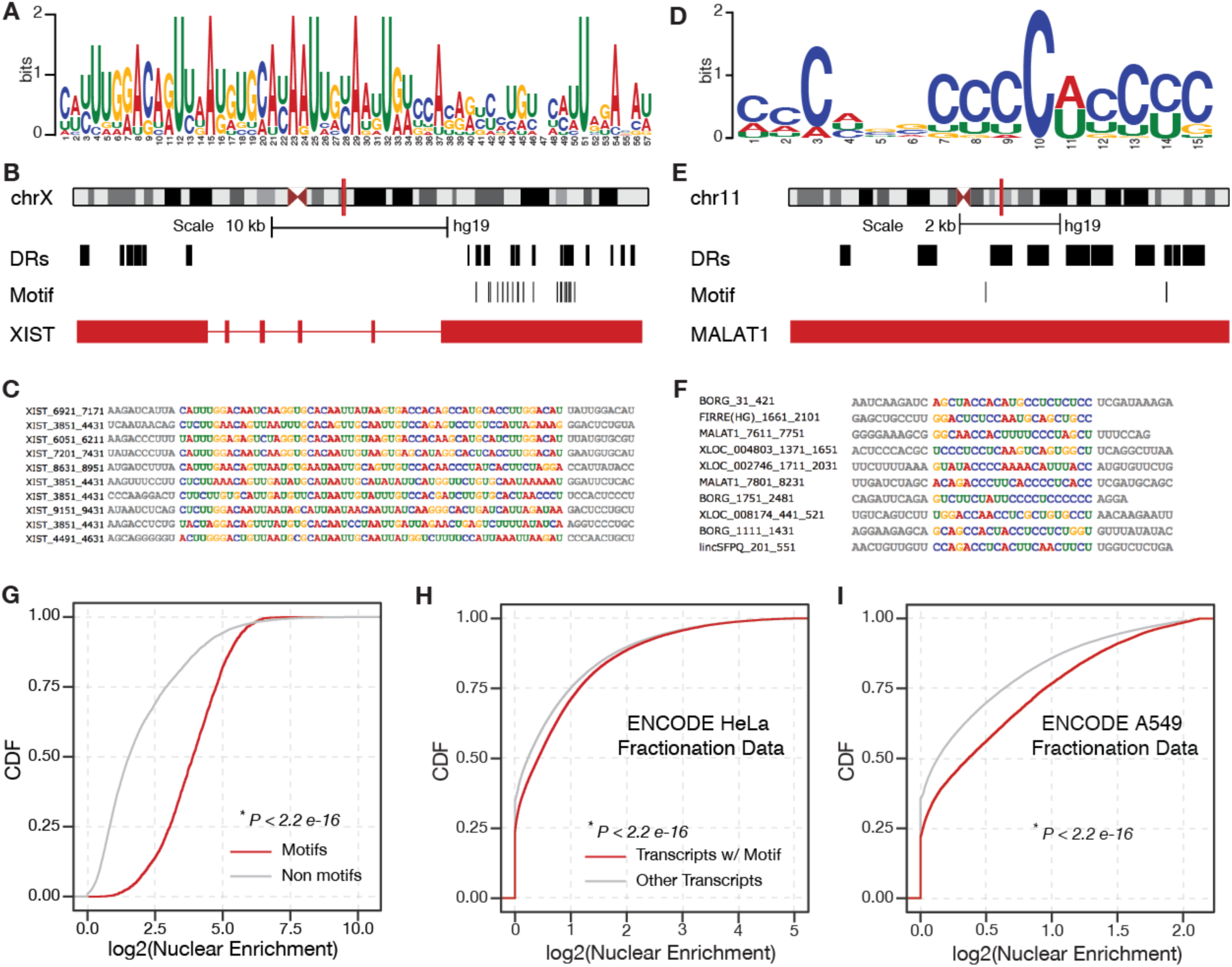
Motifs enriched in lncRNA nuclear enrichment signals. **A**. Position Weight Matrix (PWM) for a novel 57 nt motif enriched within the DRs of lncRNA *XIST*, discovered using MEME^28^. E-value < 0.05 **B**. Occurrences of this motif throughout the *XIST* locus. **C**. Multiple sequence alignment of the incidences of this *XIST* motif (*colored nucleotides*) within Differential Regions. Adjoining sequences are colored in gray. **D**. PWM for a novel C-rich 15 nt motif enriched within the DR’s of 21 different lncRNAs, discovered using MEME. E-value < 0.05 **E**. The occurrences of this motif throughout the *MALAT1* locus. **F**. Multiple sequence alignment of different instances of this motif (*colored nucleotides)*, as they appear in the Differential Regions of the indicated lncRNAs. **G.** Oligos bearing the novel motifs described in **A-F** and Extended Data Figure 4 are significantly enriched in nuclear fractions, relative to all other oligos in the MPRA pool. *P*-value: Mann Whitney Test. **H-I**. Novel nuclear enrichment motifs influence the localization of endogenous human transcripts. CDF plot comparing the nuclear enrichment of all human transcripts with at least one occurrence of our discovered motifs, relative to all other transcripts, in HeLa and A549 cells^30^. *P*-value: Mann Whitney Test.

We independently tested if these motifs are sufficient for nuclear localization using smRNA-FISH. Briefly, we appended the consensus motif sequences identified by our MPRA to the 3’ end of the cytosolic fsSox2 reporter and electroporated these constructs in HeLa cells ^9^. We then performed smRNA-FISH^31^ and did a double blinded quantification of the signals in more than 300 nuclei for each electroporated construct using StarSearch^31^ (*Methods*). We observed that ~30 % of *fsSox2* transcripts localized in the nucleus but appending the repetitive *XIST* motif (Motif 1) slightly increased nuclear localization to ~40% (Figure 4; *P* = 0.03, Mann-Whitney Test). Appending the C-rich motif (Motif 2) did not significantly affect the localization of *fsSox2* (Figure 4). These results suggest that small motifs could exhibit a weak effect of RNA nuclear enrichment, but are insufficient for localization.

**Figure 4.**
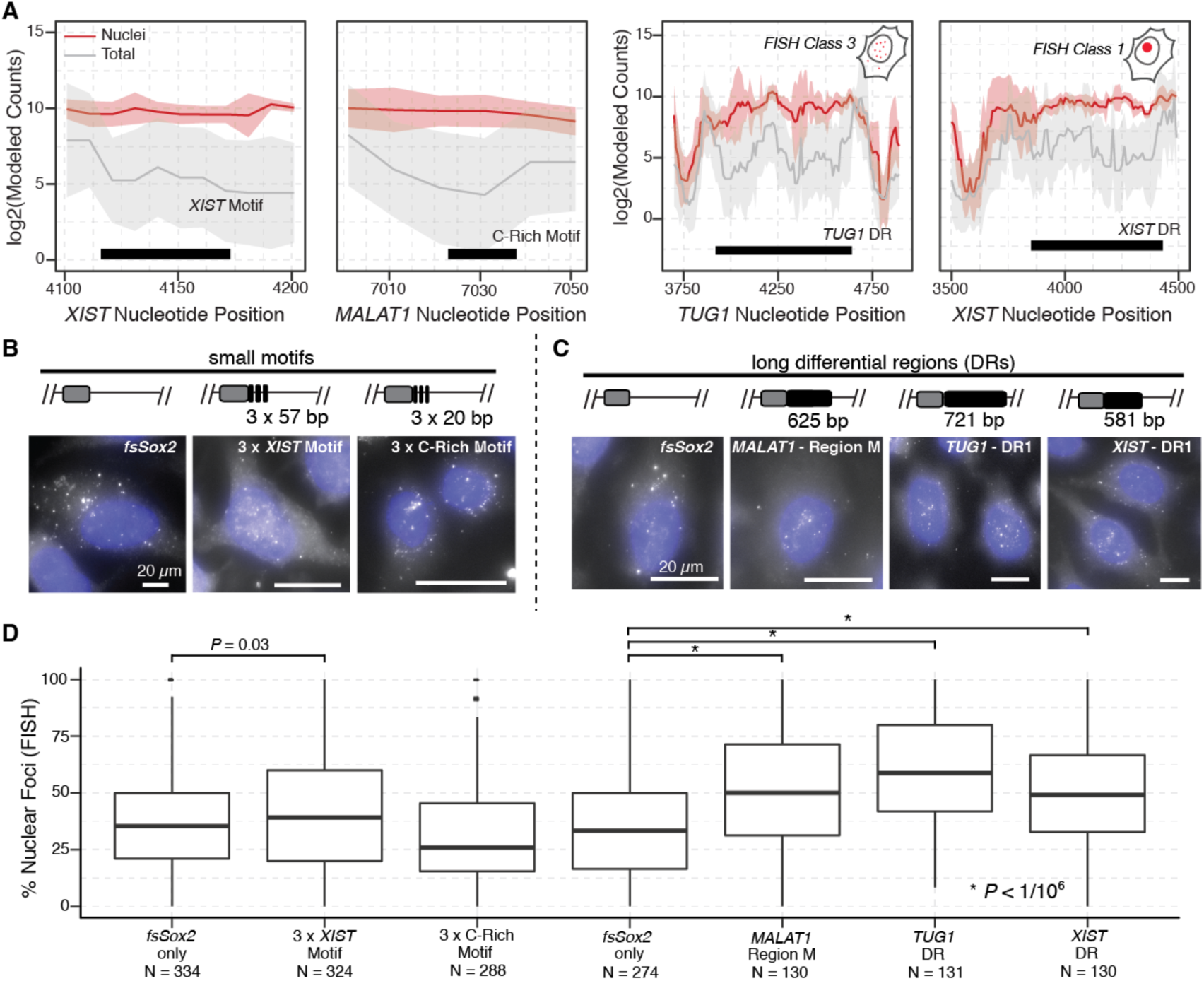
Differential Regions are sufficient to redirect RNA subcellular localization A-B. Representative *XIST* and C-Rich motif regions and novel Differential Regions from lncRNAs *TUG1* and *XIST* that are examined in **B–D.** Data depicted as in Figure 1C. **B-C.** Experimental overview of single-molecule RNA FISH (smRNA-FISH) experiments. Sox2 reporter constructs fused to individual motifs are transiently expressed in HeLa cells, and the resulting fusion transcripts are imaged using a common probe set targeting fsSox2^9, 31^. Representative smRNA-FISH images demonstrating the behavior of (*left*) the unmodified fsSox2 reporter, (*middle*) the reporter fused to three tandem instance of the XIST-derived motif (“Motif1”), and (*right*) the reporter fused to three tandem instances of the C-rich motif (“Motif2”). Scale bars are the same for all images. Blue: Hoechst 33342 Representative smRNA-FISH images of HeLa cells transiently expressing the indicated Sox2 reporter constructs: unmodified fsSox2, MALAT1 Region M (*second from left*), TUG1 Differential Region (*second from right*) and XIST Differential Region (*right most*). Data were collected using the experimental scheme outlined above. Scale bars are the same for all images. **E**. Quantification of the apparent nuclear localization of Sox2 reporter constructs, fused to the indicated Motifs, as observed using smRNA-FISH (*Methods*) *P*-value: Mann Whitney Test.

Since we observed only a small effect for a short motif like the *XIST* motif to affect nuclear enrichment, we next asked next whether longer regions identified by our MPRA would show a stronger effect. To this end we generated multiple *fsSox2*:DR constructs (DRs: *MALAT1*, *TUG1*, *XIST*) and compared their subcellular localization to the native *fsSox2*transcript by smRNA-FISH. We found that *MALAT1* “Region M” significantly increased nuclear enrichment of *fsSox2* (Figure 4; *P* < 1/10^6^, Mann-Whitney Test). Similarly, a novel *TUG1* DR identified by our MPRA, as well as *the XIST* DR, which harbors the *XIST* motif, showed also nuclear enrichment of fsSox2 (Figure 4; *P* < 1/10^6^, Mann-Whitney Test; *Methods*). Thus, the longer DRs identified in our MPRA are sufficient to significantly change the nuclear enrichment of a cytosolic transcript where as shorter motifs could not.

Collectively, our study has several implications. First, we have demonstrated a new functional MPRA which can identify longer nuclear enrichment sequences by computationally stitching short (110 bp) oligonucleotides together. Second, we have discovered motifs common to many DRs that tend to be nuclear enriched. However, these small motifs exhibit only a mild propensity for nuclear enrichment when tested independently. Conversely, longer DRs were sufficient to change the nuclear enrichment of a cytosolic reporter. While this manuscript was in preparation, a C-rich motif similar to that identified by our MPRA was also found by other investigators and functionally tested by mutation and protein binding preferences (Igor Ulitsky – personal communication). Third, many DRs identified in our study did not harbor any motif and many lncRNAs harbored multiple DRs.

Taken together, these results indicate that there does not appear to be a small universal sequence motif that is sufficient for nuclear enrichment. Rather, we propose that multiple unique sequences co-occurring within a longer structured region are responsible for nuclear enrichment for each lncRNA. While additional studies will need to confirm this prediction, our study provides an important initial map and a systematic, unbiased framework to explore RNA nuclear enrichment signals.

## Methods

### Oligo Pool Design

We designed 153-mer oligonucleotides to contain, in order, the 16-nt universal primer site ACTGGCCGCTTCACTG, a 110-nt variable sequence, a 10-nt unique barcode sequence and the 17-nt universal primer site AGATCGGAAGAGCGTCG. The unique barcodes were designed as described previously while the variable sequences were obtained by tiling lncRNA sequences. The resulting oligonucleotide libraries were synthesized by Broad Technology Labs.

### ePCR amplification of oligopool

The synthesized oligopool was amplified by emulsion-PCR (ePCR, Micellula DNA Emulsion & Purification Kit, Chimerx), according to the manufacturers’ instructions. The e-PCR primers where designed to add the Age I / Not I restriction sites to the synthesized oligos for subsequent cloning (Age I primer: AATAATACCGGTACTGGCCGCTTCACTG; Not I primer: GAGGCCGCG GCCGCCGACGCTCTTCCGATCT). To determine the oligos representation of the ePCR amplified oligo pool (based on the unique 3’ barcode of each oligo), 1 ng of the amplified oligo pool was used as input for library preparation (see below) and sequenced on a MiSeq (SR, Illumina).

### Cloning

A minCMV promoter (5’-TAGGCGTGTACGGTGGGAGGCCTATATAAGCAGAGCTCGTTTAGTGAACCGTCAGATCGC-3’) was cloned upstream of fsSox2^9^. The ePCR-amplified oligopool and the identified motifs and candidate regions were digested with Age I / Not I and inserted 3’ of fsSox2. For MPRA-cloning, the ligation reaction (100 ng backbone + 4 × molar excess of oligopool) was transformed into 10 × DH5α tubes (ThermoScientific). A total of 20 ampicillin LB plates were inoculated with the 10 transformation reactions and incubated overnight at 37°C. All bacterial colonies were then scraped in 5 ml of LB per plate and pooled, and the plasmids were purified with the endotoxin-free Qiagen Plasmid Plus Maxi kit (Qiagen). The cloned oligopool was then sequenced on the MiSeq to determine the oligo representation as described above.

### Cell fractionation

HeLa nuclear and cytoplasmic fractions were isolated as previously described^9^. The success of the fractionations (Extended Data Figure 2B) was confirmed by qRT-PCR of the nuclear ncRNA NEAT1 and the cytoplasmic ncRNA SNHG5 in RNA isolated (see below) from whole cells, the pelleted nuclei, and from the cytoplasmic fractions.

### RNA extraction and qRT-PCR

RNA was isolated by TRIzol (ThermoScientific) – chloroform extraction, followed by isopropanol precipitation, according to standard procedures. 2 *μ*g of BioAnalyzer-validated RNA were digested with recombinant DNase-I (2.77 U/*μ*l, Worthington #LS006353) at 37°C for 30 min, followed by heat-inactivation at 75°C for 10 min. Reverse transcription was performed with SuperScript III cDNA synthesis kit (ThermoScientific). Quantitative RT-PCR was performed using the FastStart Universal SYBR Green Master mix (Roche) on an ABI 7900. Primers were: NEAT1 forward TGATGCCACAACGCAGATTG, reverse GCAAACAGGTGGGTAGGTGA, and SNHG5 forward GTGGACGAGTAGCCAGTGAA, reverse GCCTCTATCAATGGGCAGACA. After processing the raw data by qPCR Miner^32^, the efficiency of each primer set was used to calculate the relative initial concentration of each gene. The relative expression in the nuclear and cytoplasmic fractions was then calculated by normalization to that in the whole cell.

### Library preparation

Sequencing libraries were prepared by PCR amplification using PfuUltra II Fusion DNA polymerase (Agilent #600672) and primers designed to anneal to the universal primer site flanking the oligos and to add sequencing index barcode for multiplexing: forward caagcagaagacggcatacgagatCGTGATgtgactggagttcagacgtgtgctcttccgatctACTGGCCGCTTCACT G, reverse AATGATACGGCGACCACCGAGATCTACACTCTTTCCCTACACGACGCTCTTCCG ATCT (capital letters indicate (1) the index for the library and (2) the region complementary to the universal primer site). PCR amplification (initial denaturation 95°C – 2 min; cycling 95°C – 30 secs, 55°C – 30 secs, 72°C – 30 sec; final extension 72°C – 10 min) was carried out for 30 cycles followed by triple 0.6×, 1.6×, and 1× SPRI beads (Agencourt AMPure XP, Beckman Coulter) cleanup. The quality and molarity of the libraries was evaluated by BioAnalyzer and the samples were sequenced in a pool of 6 on the Illumina HiSeq2500, full flow cell, single-read 100 bp. To ensure the transfection was successful, we required that at least 70% of the oligo pool was represented back (i.e. had a count of at least one) in the sequencing sample. (Extended Data Figure 2, 3 and 4)

## Analyzing MPRA Data

### Read Mapping and Obtaining Counts Table

To find a unique mapping location for the read, we ensured an exact match between the first 10 read nucleotides and a unique oligo barcode. To ensure that the correct oligo was identified using this barcode match, we allowed only 2 mismatches between the remaining 65 nts of the read sequence and the upstream oligo sequence corresponding to the unique barcode (Extended Data Figure 1A). The resulting counts for each oligo in every sample (6 Nuclei and 6 Total) were compiled in a counts table (Extended Data Figure 1A).

### Normalizing the counts table

The counts table was normalized using a library size correction in order to facilitate comparing counts across samples with different sequencing depths. The library size was calculated as the total number of reads in each sample.

### Modeling Nucleotide Counts from Oligo Counts

The counts of a particular nucleotide were modeled by taking the median of counts for every oligo tiling the nucleotide (Extended Data Figure 1B). We tried other methods to model nucleotide such as taking the sum of the counts of all oligos tiling the given nucleotide and a probabilistic graphical model as used recently^15^ but the simple and intuitive median approach yielded comparable results. Since the offset between subsequent oligos was usually 10 nucleotides, we obtained nucleotide counts also at a 10 nucleotide resolution. The resulting modeled nucleotide counts table (Extended Datat Table 2) was used to infer differential regions.

### Inferring Differential Regions from Modeled Nucleotide Counts

There are 2 main steps in inferring differential regions from modeled nucleotide counts – (i). Identifying potential candidate regions and (ii). Assigning a p-value for each potential candidate region (Extended Data Figure 1C). We identified potential candidate regions by calculating the median of the difference between nuclear counts and total counts across all 6 replicates at each nucleotide and then grouping together neighboring points that exceeded a threshold, as described previously^25^. We then defined a summary statistics for each region based on the differences between nuclear and total counts of each nucleotide in the region as well as the trend of these counts. To assess the uncertainty of this procedure we generated a list of global null candidates by shuffling the sample labels and computed a summary statistic for these regions to form a null distribution. Then we ranked each potential candidate region by comparing their respective summary statistic to the null distribution to obtain an empirical p-value. The p-values were converted to q-values using the Benjamini-Hochberg approach.

### Motif Analysis

MEME software package was used to find motifs enriched in differential regions. Specifically, we used the MEME function in the suite in the discriminative mode with DR sequences as the list of primary sequences and the other sequences in the pool as the controls. We ran MEME in different settings – OOPS and ANR - to ensure we found motifs that were repeating several times in a given DR and those only occurring once.

### K-mer Enrichment

If sequence preferences are driven by more general sequence composition preferences that cannot be so easily represented by regular expression or position weight matrix motif models, then nuclear enrichment of DRs may be more effectively modeled by considering all k-mers. To this end, we performed a regression to assign weight coefficients to all k-mers for the DR sequences and non-DR sequences similar to the motif analysis using MEME as described previously. To avoid overfitting, we performed ridge regression^29^, which minimizes not only the distance between model predictions and actual values but also the magnitude of the weights. We chose the alpha parameter that varies the emphasis of these two competing objectives by evaluating fivefold cross-validated mean squared error over a parameter grid.

### Conservation Analysis

The phastCons and phyloP scores for the whole genome were downloaded from UCSC genome browser. We extracted these scores for the DRs and shuffled control regions using a custom script. In order to account for natural conservation differences between lncRNAs and mRNAs as well as among different lncRNAs, the control regions were obtained by shuffling the DR sequences using shuffleBed but ensuring the new regions fell within exons of the lncRNAs the DRs were from. Finally, the scores were compared between DR and non-DR regions using the Mann-Whitney test.

### ENCODE Fractionation RNA-Seq

We downloaded the raw RNA-Seq reads for the nucleus and cytosolic compartments from the ENCODE^30^ website. These reads were quantified using kallisto to obtain TPMs and then the nuclear/cytosolic TPMs of transcripts with the motif (found using the FIMO software) were compared to all the other transcripts.

### Single molecule RNA fluorescence *in situ* hybridization (smRNA FISH)

Briefly, 70-80% confluent 1×10^6^ HeLa [ATCC^®^ CCL-2^TM^] cells were electroporated with 2 μg of construct using the Amaxa^®^ Cell Line Nucleofector^®^ Kit R using program I-013, and cultured for 48 hours in LabTek v1 glass chambers. smRNA-FISH was performed using Biosearch Technologies Stellaris^®^ probes, as described previously (Reference). RNA probes targeting and tiling the fsSox2 exon were conjugated to Quasar 570. Nuclei were visualized with 4,6-diamidino-2-phenylindole (DAPI). Images were obtained using the Zeiss Cell Observer Live Cell microscope at the Harvard Center for Biological Imaging. For each field of view, at least 40 slices (each plane: 0.24 μm) were imaged, and z-stacks were merged with maximum intensity projections (MIP). Sox2 foci were computationally-identified using the spot counting software StarSearch. To ensure robustness, the analysis was blinded and the person counting the spots did not know the identity of the samples. For each construct, fsSox2 foci within at least 150 cells were counted in biological duplicate.

## Code availability

All the analysis in this paper was carried out using a custom package developed for the experiment called oligoGames. The package is currently hosted on GitHub – https://github.com/cshukla/oligoGames.

## Data availability

All analyzed sequence data has been deposited in NCBI GEO under accession GSE98828. References

**Supplementary Information** is available in the online version of the paper

## Acknowledgements

The authors would like to thank Doug Richardson and Sven Terclavers at Harvard Center for Biological Imaging for assistance with imaging, Ezgi Haceysuleyman for advice on smRNA-FISH and Bauer Sequencing Facility at Harvard University for assistance with the sequencing. CJS would like to acknowledge Alejandro Reyes for advice on writing the manuscript and analyzing the data. The authors would like to thank everyone in the Rinn and Irizarry lab for their advice and insightful comments throughout this work. This work was supported by NIH grants R01GM083084 and R01HG005220 to RAI as well as NIH grants U01DA040612-01 and P01GM099117 to JLR.

## Author Contributions

These authors contributed equally to this work.

Philipp G Maass, John L Rinn

## Author Information

## Tables

**Extended Data Table 1:**
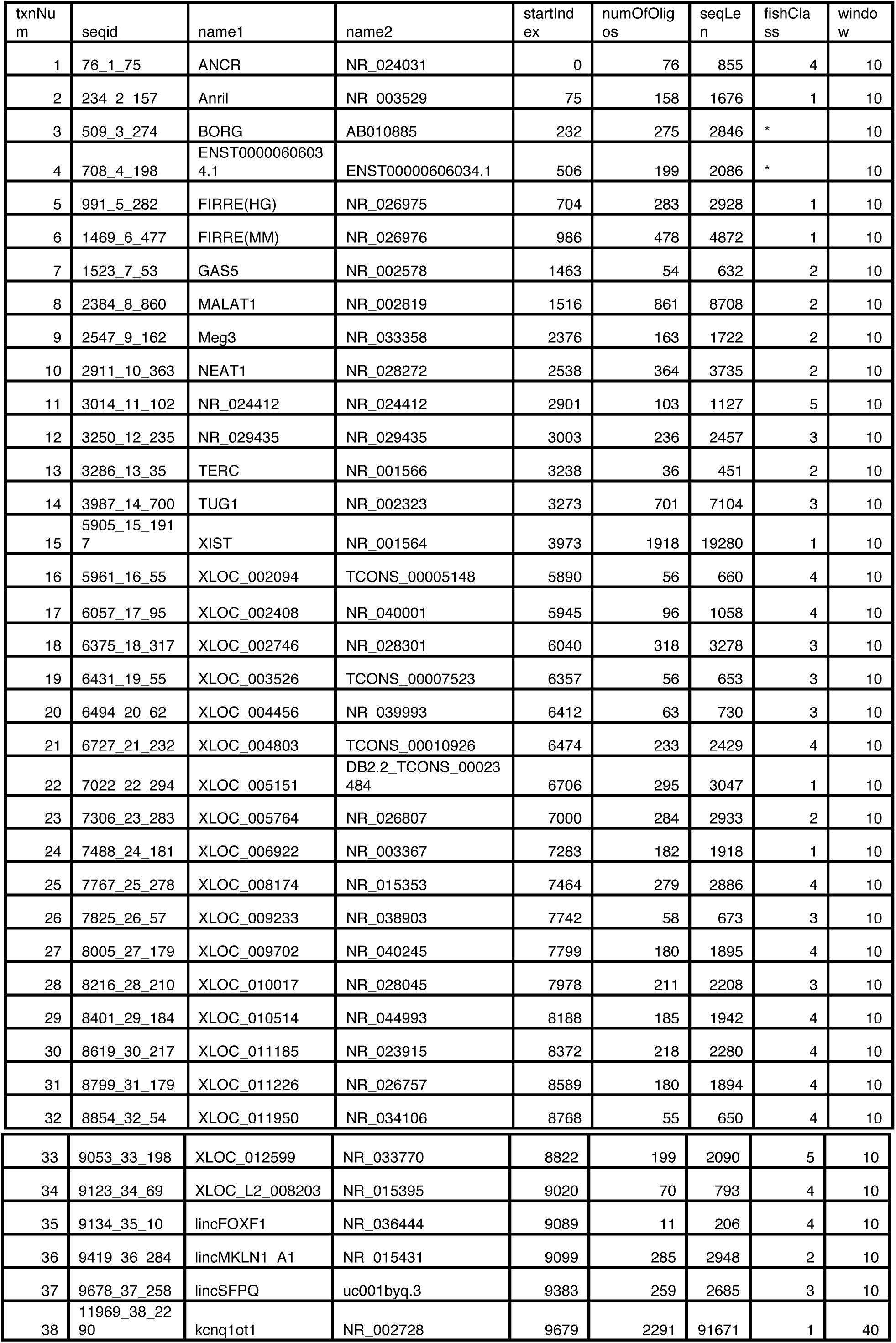
A table describing the meta data of the oligo pool used in this work.

**Extended Data Table 2:**
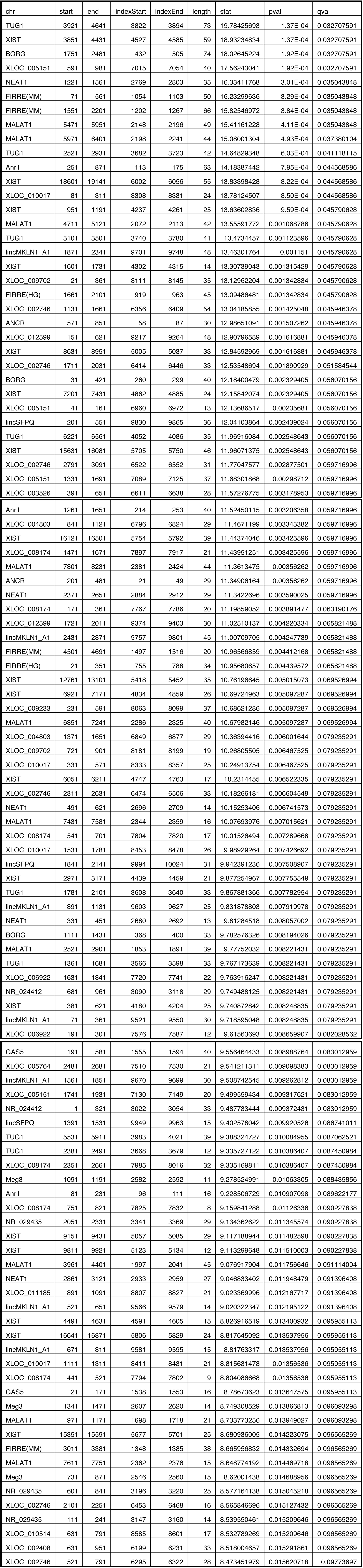
A table describing the 109 DRs discovered in this work.

| chr | start | end | indexStart | indexEnd | length | stat | pval | qval |
| --- | --- | --- | --- | --- | --- | --- | --- | --- |
| TUG1 | 3921 | 4641 | 3822 | 3894 | 73 | 19.78425693 | 1.37E-04 | 0.032707591 |
| XIST | 3851 | 4431 | 4527 | 4585 | 59 | 18.93234834 | 1.37E-04 | 0.032707591 |
| BORG | 1751 | 2481 | 432 | 505 | 74 | 18.02645224 | 1.92E-04 | 0.032707591 |
| XLOC_005151 | 591 | 981 | 7015 | 7054 | 40 | 17.56243041 | 1.92E-04 | 0.032707591 |
| NEAT1 | 1221 | 1561 | 2769 | 2803 | 35 | 16.33411768 | 3.01E-04 | 0.035043848 |
| FIRRE(MM) | 71 | 561 | 1054 | 1103 | 50 | 16.23299636 | 3.29E-04 | 0.035043848 |
| FIRRE(MM) | 1551 | 2201 | 1202 | 1267 | 66 | 15.82546972 | 3.84E-04 | 0.035043848 |
| MALAT1 | 5471 | 5951 | 2148 | 2196 | 49 | 15.41161228 | 4.11E-04 | 0.035043848 |
| MALAT1 | 5971 | 6401 | 2198 | 2241 | 44 | 15.08001304 | 4.93E-04 | 0.037380104 |
| TUG1 | 2521 | 2931 | 3682 | 3723 | 42 | 14.64829348 | 6.03E-04 | 0.041118115 |
| Anril | 251 | 871 | 113 | 175 | 63 | 14.18387442 | 7.95E-04 | 0.044568586 |
| XIST | 18601 | 19141 | 6002 | 6056 | 55 | 13.83398428 | 8.22E-04 | 0.044568586 |
| XLOC_010017 | 81 | 311 | 8308 | 8331 | 24 | 13.78124507 | 8.50E-04 | 0.044568586 |
| XIST | 951 | 1191 | 4237 | 4261 | 25 | 13.63602836 | 9.59E-04 | 0.045790628 |
| MALAT1 | 4711 | 5121 | 2072 | 2113 | 42 | 13.55591772 | 0.001068786 | 0.045790628 |
| TUG1 | 3101 | 3501 | 3740 | 3780 | 41 | 13.4734457 | 0.001123596 | 0.045790628 |
| lincMKLN1_A1 | 1871 | 2341 | 9701 | 9748 | 48 | 13.46301764 | 0.001151 | 0.045790628 |
| XIST | 1601 | 1731 | 4302 | 4315 | 14 | 13.30739043 | 0.001315429 | 0.045790628 |
| XLOC_009702 | 21 | 361 | 8111 | 8145 | 35 | 13.12962204 | 0.001342834 | 0.045790628 |
| FIRRE(HG) | 1661 | 2101 | 919 | 963 | 45 | 13.09486481 | 0.001342834 | 0.045790628 |
| XLOC_002746 | 1131 | 1661 | 6356 | 6409 | 54 | 13.04185855 | 0.001425048 | 0.045946378 |
| ANCR | 571 | 851 | 58 | 87 | 30 | 12.98651091 | 0.001507262 | 0.045946378 |
| XLOC_012599 | 151 | 621 | 9217 | 9264 | 48 | 12.90796589 | 0.001616881 | 0.045946378 |
| XIST | 8631 | 8951 | 5005 | 5037 | 33 | 12.84592969 | 0.001616881 | 0.045946378 |
| XLOC_002746 | 1711 | 2031 | 6414 | 6446 | 33 | 12.53548694 | 0.001890929 | 0.051584544 |
| BORG | 31 | 421 | 260 | 299 | 40 | 12.18400479 | 0.002329405 | 0.056070156 |
| XIST | 7201 | 7431 | 4862 | 4885 | 24 | 12.15842074 | 0.002329405 | 0.056070156 |
| XLOC_005151 | 41 | 161 | 6960 | 6972 | 13 | 12.13686517 | 0.00235681 | 0.056070156 |
| lincSFPQ | 201 | 551 | 9830 | 9865 | 36 | 12.04103864 | 0.002439024 | 0.056070156 |
| TUG1 | 6221 | 6561 | 4052 | 4086 | 35 | 11.96916084 | 0.002548643 | 0.056070156 |
| XIST | 15631 | 16081 | 5705 | 5750 | 46 | 11.96071375 | 0.002548643 | 0.056070156 |
| XLOC_002746 | 2791 | 3091 | 6522 | 6552 | 31 | 11.77047577 | 0.002877501 | 0.059716996 |
| XLOC_005151 | 1331 | 1691 | 7089 | 7125 | 37 | 11.68301868 | 0.00298712 | 0.059716996 |
| XLOC_003526 | 391 | 651 | 6611 | 6638 | 28 | 11.57276775 | 0.003178953 | 0.059716996 |
| Anril | 1261 | 1651 | 214 | 253 | 40 | 11.52450115 | 0.003206358 | 0.059716996 |
| XLOC_004803 | 841 | 1121 | 6796 | 6824 | 29 | 11.4671199 | 0.003343382 | 0.059716996 |
| XIST | 16121 | 16501 | 5754 | 5792 | 39 | 11.44374046 | 0.003425596 | 0.059716996 |
| XLOC_008174 | 1471 | 1671 | 7897 | 7917 | 21 | 11.43951251 | 0.003425596 | 0.059716996 |
| MALAT1 | 7801 | 8231 | 2381 | 2424 | 44 | 11.3613475 | 0.00356262 | 0.059716996 |
| ANCR | 201 | 481 | 21 | 49 | 29 | 11.34906164 | 0.00356262 | 0.059716996 |
| NEAT1 | 2371 | 2651 | 2884 | 2912 | 29 | 11.3422696 | 0.003590025 | 0.059716996 |
| XLOC_008174 | 171 | 361 | 7767 | 7786 | 20 | 11.19859052 | 0.003891477 | 0.063190176 |
| XLOC_012599 | 1721 | 2011 | 9374 | 9403 | 30 | 11.02510137 | 0.004220334 | 0.065821488 |
| lincMKLN1_A1 | 2431 | 2871 | 9757 | 9801 | 45 | 11.00709705 | 0.004247739 | 0.065821488 |
| FIRRE(MM) | 4501 | 4691 | 1497 | 1516 | 20 | 10.96566859 | 0.004412168 | 0.065821488 |
| FIRRE(HG) | 21 | 351 | 755 | 788 | 34 | 10.95680657 | 0.004439572 | 0.065821488 |
| XIST | 12761 | 13101 | 5418 | 5452 | 35 | 10.76196645 | 0.005015073 | 0.069526994 |
| XIST | 6921 | 7171 | 4834 | 4859 | 26 | 10.69724963 | 0.005097287 | 0.069526994 |
| XLOC_009233 | 231 | 591 | 8063 | 8099 | 37 | 10.68621286 | 0.005097287 | 0.069526994 |
| MALAT1 | 6851 | 7241 | 2286 | 2325 | 40 | 10.67982146 | 0.005097287 | 0.069526994 |
| XLOC_004803 | 1371 | 1651 | 6849 | 6877 | 29 | 10.36394416 | 0.006001644 | 0.079235291 |
| XLOC_009702 | 721 | 901 | 8181 | 8199 | 19 | 10.26805505 | 0.006467525 | 0.079235291 |
| XLOC_010017 | 331 | 571 | 8333 | 8357 | 25 | 10.24913754 | 0.006467525 | 0.079235291 |
| XIST | 6051 | 6211 | 4747 | 4763 | 17 | 10.2314455 | 0.006522335 | 0.079235291 |
| XLOC_002746 | 2311 | 2631 | 6474 | 6506 | 33 | 10.18266181 | 0.006604549 | 0.079235291 |
| NEAT1 | 491 | 621 | 2696 | 2709 | 14 | 10.15253406 | 0.006741573 | 0.079235291 |
| MALAT1 | 7431 | 7581 | 2344 | 2359 | 16 | 10.07693976 | 0.007015621 | 0.079235291 |
| XLOC_008174 | 541 | 701 | 7804 | 7820 | 17 | 10.01526494 | 0.007289668 | 0.079235291 |
| XLOC_010017 | 1531 | 1781 | 8453 | 8478 | 26 | 9.98929264 | 0.007426692 | 0.079235291 |
| lincSFPQ | 1841 | 2141 | 9994 | 10024 | 31 | 9.942391236 | 0.007508907 | 0.079235291 |
| XIST | 2971 | 3171 | 4439 | 4459 | 21 | 9.877254967 | 0.007755549 | 0.079235291 |
| TUG1 | 1781 | 2101 | 3608 | 3640 | 33 | 9.867881366 | 0.007782954 | 0.079235291 |
| lincMKLN1_A1 | 891 | 1131 | 9603 | 9627 | 25 | 9.831878803 | 0.007919978 | 0.079235291 |
| NEAT1 | 331 | 451 | 2680 | 2692 | 13 | 9.81284518 | 0.008057002 | 0.079235291 |
| BORG | 1111 | 1431 | 368 | 400 | 33 | 9.782576326 | 0.008194026 | 0.079235291 |
| MALAT1 | 2521 | 2901 | 1853 | 1891 | 39 | 9.77752032 | 0.008221431 | 0.079235291 |
| TUG1 | 1361 | 1681 | 3566 | 3598 | 33 | 9.767173639 | 0.008221431 | 0.079235291 |
| XLOC_006922 | 1631 | 1841 | 7720 | 7741 | 22 | 9.763916247 | 0.008221431 | 0.079235291 |
| NR_024412 | 681 | 961 | 3090 | 3118 | 29 | 9.749488125 | 0.008221431 | 0.079235291 |
| XIST | 381 | 621 | 4180 | 4204 | 25 | 9.740872842 | 0.008248835 | 0.079235291 |
| lincMKLN1_A1 | 71 | 361 | 9521 | 9550 | 30 | 9.718595048 | 0.008248835 | 0.079235291 |
| XLOC_006922 | 191 | 301 | 7576 | 7587 | 12 | 9.61563693 | 0.008659907 | 0.082028562 |
| GAS5 | 191 | 581 | 1555 | 1594 | 40 | 9.556464433 | 0.008988764 | 0.083012959 |
| XLOC_005764 | 2481 | 2681 | 7510 | 7530 | 21 | 9.541211311 | 0.009098383 | 0.083012959 |
| lincMKLN1_A1 | 1561 | 1851 | 9670 | 9699 | 30 | 9.508742545 | 0.009262812 | 0.083012959 |
| XLOC_005151 | 1741 | 1931 | 7130 | 7149 | 20 | 9.499559434 | 0.009317621 | 0.083012959 |
| NR_024412 | 1 | 321 | 3022 | 3054 | 33 | 9.487733444 | 0.009372431 | 0.083012959 |
| lincSFPQ | 1391 | 1531 | 9949 | 9963 | 15 | 9.402578042 | 0.009920526 | 0.086741011 |
| TUG1 | 5531 | 5911 | 3983 | 4021 | 39 | 9.388324727 | 0.010084955 | 0.087062521 |
| TUG1 | 2381 | 2491 | 3668 | 3679 | 12 | 9.335727122 | 0.010386407 | 0.087450984 |
| XLOC_008174 | 2351 | 2661 | 7985 | 8016 | 32 | 9.335169811 | 0.010386407 | 0.087450984 |
| Meg3 | 1091 | 1191 | 2582 | 2592 | 11 | 9.278524991 | 0.01063305 | 0.088435856 |
| Anril | 81 | 231 | 96 | 111 | 16 | 9.228506729 | 0.010907098 | 0.089622177 |
| XLOC_008174 | 751 | 821 | 7825 | 7832 | 8 | 9.159841288 | 0.01126336 | 0.090227838 |
| NR_029435 | 2051 | 2331 | 3341 | 3369 | 29 | 9.134362622 | 0.011345574 | 0.090227838 |
| XIST | 9151 | 9431 | 5057 | 5085 | 29 | 9.117188944 | 0.011482598 | 0.090227838 |
| XIST | 9811 | 9921 | 5123 | 5134 | 12 | 9.113299648 | 0.011510003 | 0.090227838 |
| MALAT1 | 3961 | 4401 | 1997 | 2041 | 45 | 9.076917904 | 0.011756646 | 0.091114004 |
| NEAT1 | 2861 | 3121 | 2933 | 2959 | 27 | 9.046833402 | 0.011948479 | 0.091396408 |
| XLOC_011185 | 891 | 1091 | 8807 | 8827 | 21 | 9.023369996 | 0.012167717 | 0.091396408 |
| lincMKLN1_A1 | 521 | 651 | 9566 | 9579 | 14 | 9.020322347 | 0.012195122 | 0.091396408 |
| XIST | 4491 | 4631 | 4591 | 4605 | 15 | 8.826916519 | 0.013400932 | 0.095955113 |
| XIST | 16641 | 16871 | 5806 | 5829 | 24 | 8.817645092 | 0.013537956 | 0.095955113 |
| lincMKLN1_A1 | 671 | 811 | 9581 | 9595 | 15 | 8.81763317 | 0.013537956 | 0.095955113 |
| XLOC_010017 | 1111 | 1311 | 8411 | 8431 | 21 | 8.815631478 | 0.01356536 | 0.095955113 |
| XLOC_008174 | 441 | 521 | 7794 | 7802 | 9 | 8.804086668 | 0.01356536 | 0.095955113 |
| GAS5 | 21 | 171 | 1538 | 1553 | 16 | 8.78673623 | 0.013647575 | 0.095955113 |
| Meg3 | 1341 | 1471 | 2607 | 2620 | 14 | 8.749308529 | 0.013866813 | 0.096093298 |
| MALAT1 | 971 | 1171 | 1698 | 1718 | 21 | 8.733773256 | 0.013949027 | 0.096093298 |
| XIST | 15351 | 15591 | 5677 | 5701 | 25 | 8.680936005 | 0.014223075 | 0.096565269 |
| FIRRE(MM) | 3011 | 3381 | 1348 | 1385 | 38 | 8.665956832 | 0.014332694 | 0.096565269 |
| MALAT1 | 7611 | 7751 | 2362 | 2376 | 15 | 8.648774192 | 0.014469718 | 0.096565269 |
| Meg3 | 731 | 871 | 2546 | 2560 | 15 | 8.62001438 | 0.014688956 | 0.096565269 |
| NR_029435 | 601 | 841 | 3196 | 3220 | 25 | 8.577164138 | 0.015045218 | 0.096565269 |
| XLOC_002746 | 2101 | 2251 | 6453 | 6468 | 16 | 8.565846696 | 0.015127432 | 0.096565269 |
| NR_029435 | 111 | 241 | 3147 | 3160 | 14 | 8.539550461 | 0.015209646 | 0.096565269 |
| XLOC_010514 | 631 | 791 | 8585 | 8601 | 17 | 8.532789269 | 0.015209646 | 0.096565269 |
| XLOC_002408 | 631 | 951 | 6199 | 6231 | 33 | 8.518004657 | 0.015291861 | 0.096565269 |
| XLOC_002746 | 521 | 791 | 6295 | 6322 | 28 | 8.473451979 | 0.015620718 | 0.09773697 |

## Extended Data Figure Legends

**Figure 1.**
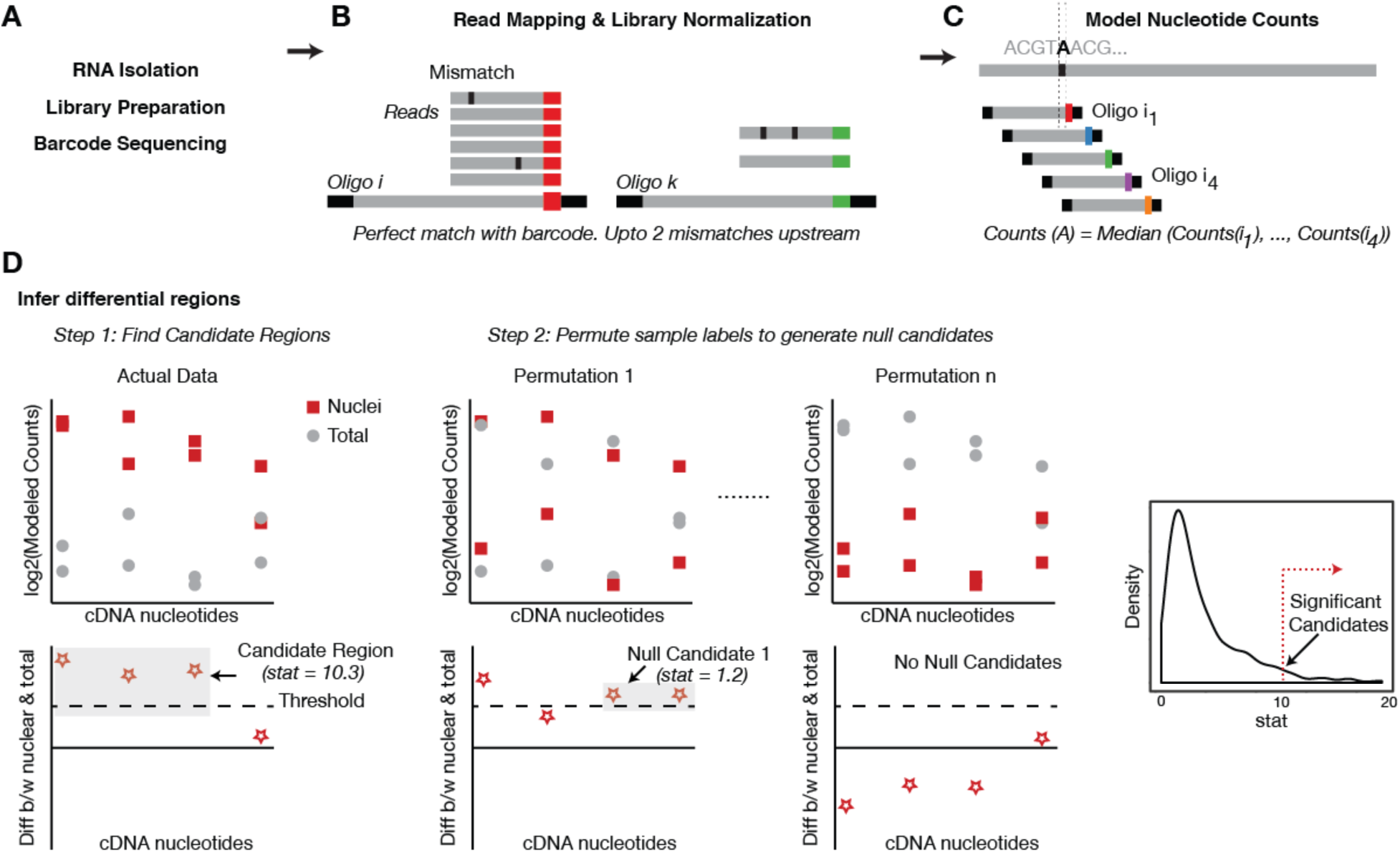
Computational pipeline to identify nuclear enrichment signals from MPRA data. **A.** Post fractionation, RNA from the nucleus and whole cell lysate is extracted. Using the universal primer sequences, the oligos are amplified in a targeted manner to make the library which is sent for sequencing. **B**. The first step in the analysis process is to map the reads back to the oligo pool. Due to the dense tiling of lncRNAs in our pool, we ensure there is a perfect match between the first 10 nucleotides of the read and the barcode sequence to ‘map’ the read. Next, we require the upstream 90 bps to only have 2 mismatches to guarantee robusteness of the mapping procedure. This step is performed by the ‘mapReads’ function in our package which gives a table of counts for each oligo as the output. This counts table is subsequently normalized for library size using the ‘normCounts’ table. We provide this normalized counts table along with the data on GEO **C**. Based on the normalized counts for each oligo, counts for each nucleotide are modeled next. As shown in the schematic, if a nucleotide ‘A’ overlaps with oligos i_1_, i_2_, i_3_ and i_4_ the counts for the nucleotide A are modeled by taking the median of counts for each of the individual oligos i_1_ – i_4_. We use the ‘modelNucCounts’ function in our package for this and get a counts table for each nucleotide in all the 12 samples (6 nuclei and 6 total) as the output (Supplementary File 2). **D**. Using the nucleotide counts table, we infer differential regions by 1). Finding candidate regions and assigning a summary statistic to each one of them and 2). Generating null candidates by permuting sample labels and using them to assign an empirical *P*-value to our candidate regions from Step 1. Please see Methods for more details (*Inset*) A distribution of the summary statitistic generated for the data we present in the paper – the red line shows the cutoff used to decide the ‘significant’ candidates.

**Figure 2.**
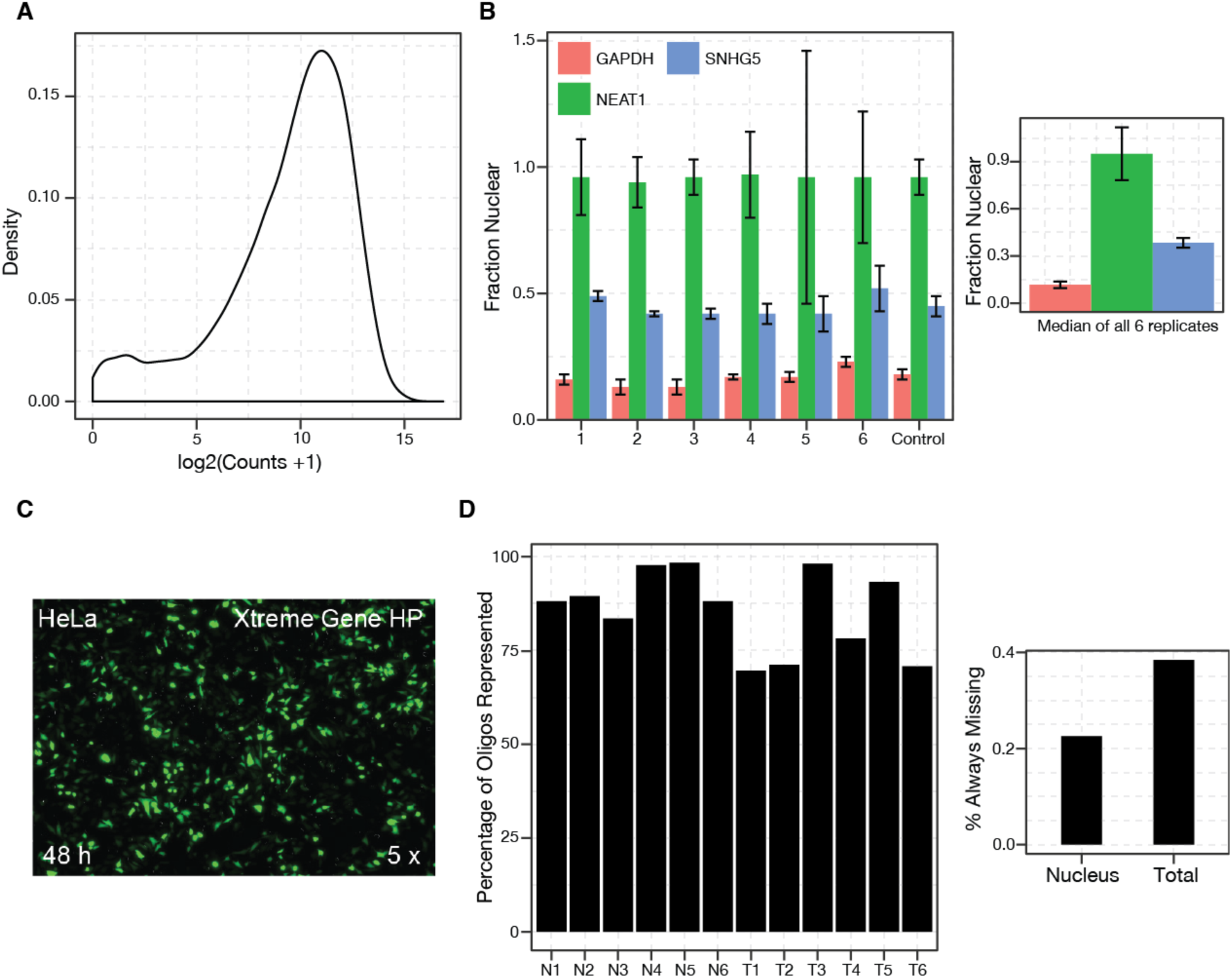
Quality Control for Various MPRA Steps. Since the MPRA has several steps, we used controls at every stage to make sure the assay was working as designed **A**. The distribution of oligo’s in our cloned plasmid pool. We see that (i). there is very little jackpotting (just a single peak showing uniform counts for several different oligos) and (ii). we have almost the entire pool represented (very small bump at zero counts). **B**. The nuclear enrichment of *NEAT1, GAPDH* and *SNHG5* as determined by qRT-PCR (*Methods*). The error bars represent standard deviation for each measurement. We see that the lncRNA *NEAT1* (green) is enriched in the nuclear fraction as expected while The ‘control’ represents the enrichment of the genes in untransfected cells (*Inset*) The median enrichment of the genes across all 6 replicates. **C**. A representative image of HeLa cells co-transfected with a GFP plasmid using the protocol outlined in Methods showing that we achieve a high transfection efficiency. **D**. The number of oligos ‘missing’ (i.e. with zero counts) from each of our 12 samples. We see that we recover >70% of our initial pool in each sample and looking across the 6 samples for nucleus and total, only 0.2% oligos (i.e. ~25 oligos) are missing from the nuclear samples and ~0.4% (i.e. ~50 oligos) are missing from the total sample.

**Figure 3.**
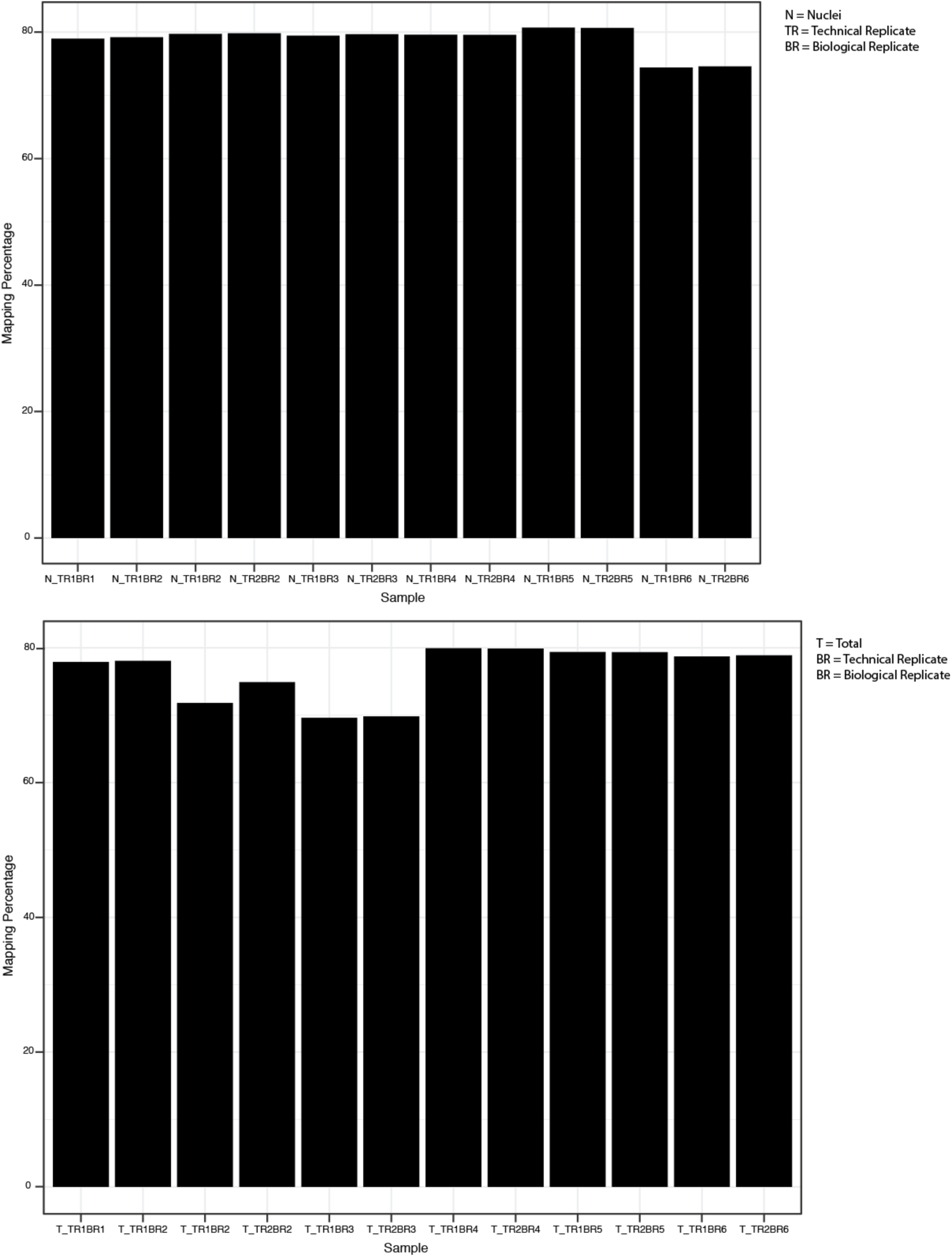
Mapping Rates for our different samples. A bar plot showing the mapping percentage for all reads of different samples from nuclear fraction (N) and total fraction (T). We show the mapping rates separately for the 2 technical replicates (TR) and each of the 6 biological replicates (BR).

**Figure 4.**
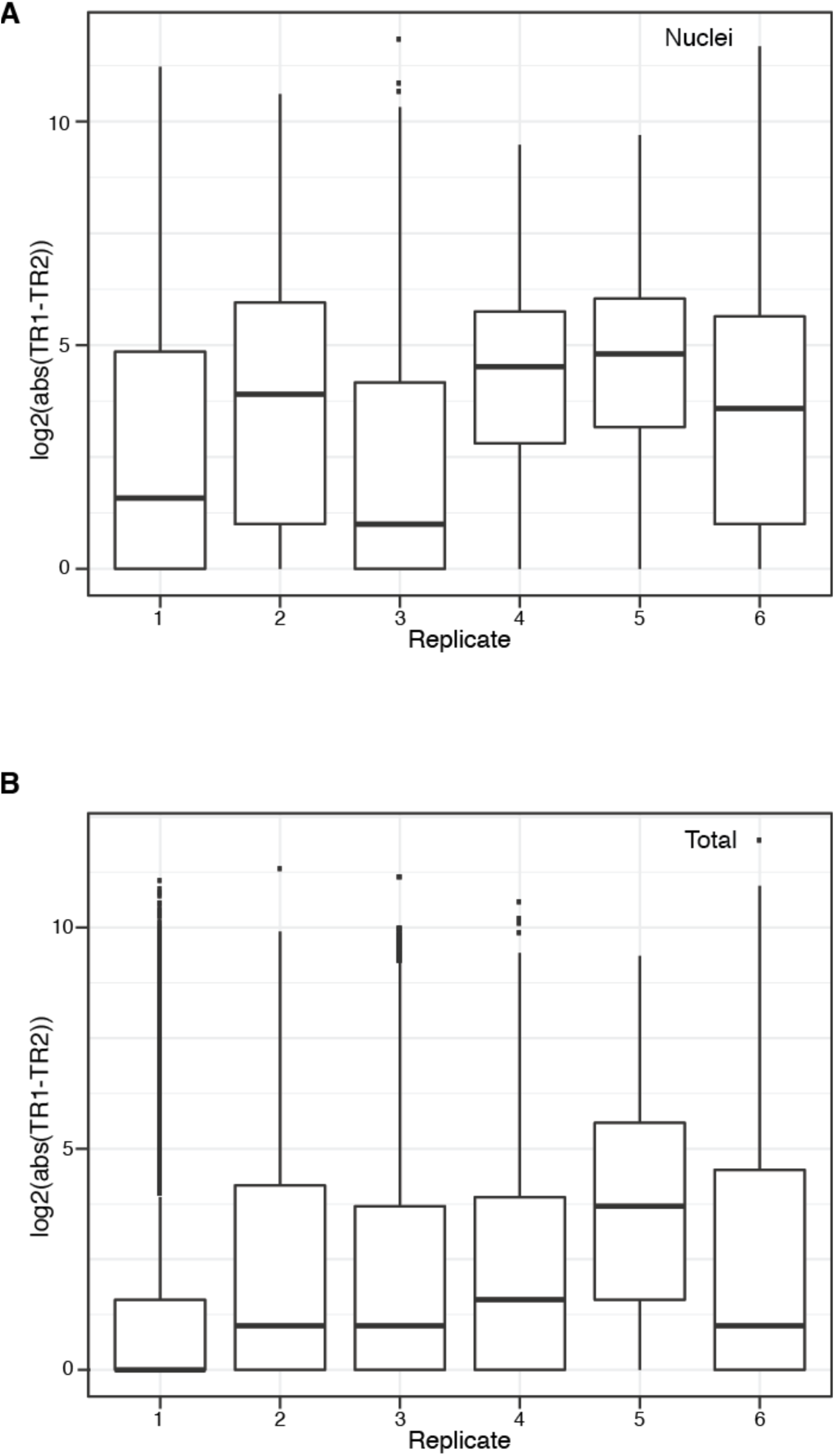
Difference between counts of technical replicates. A boxplot showing difference between counts of same oligo between the 2 technical replicates. We see that many oligos show very low difference in counts among technical replicates and thus there is very low technical variance.

**Figure 5.**
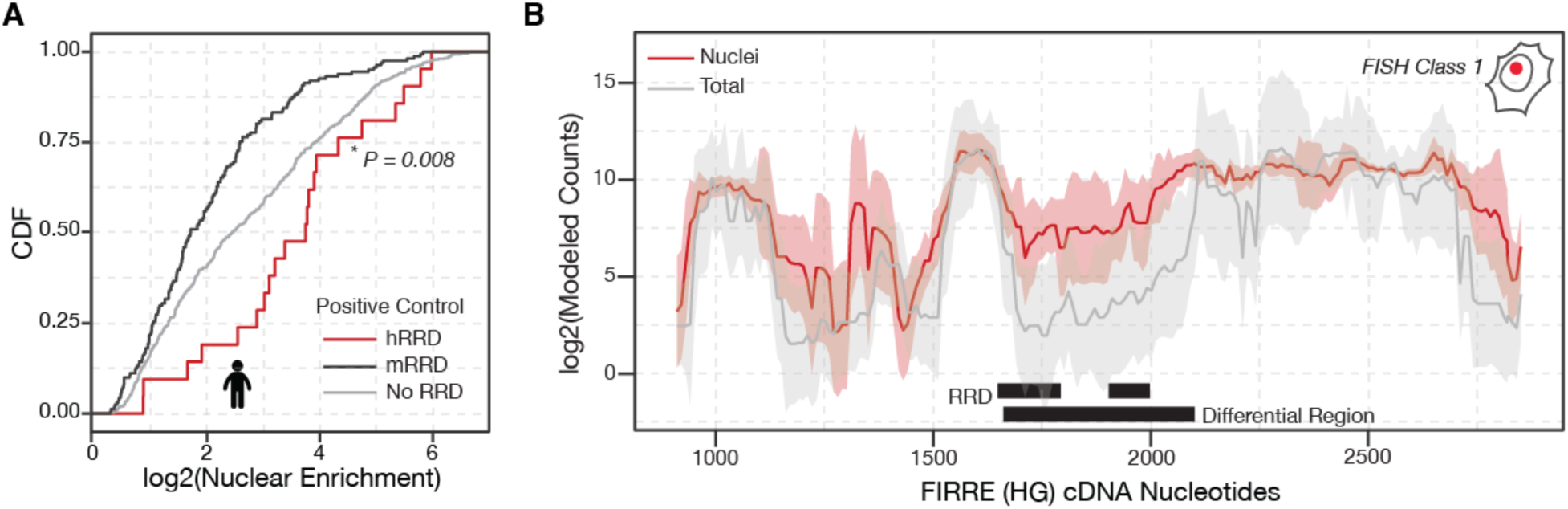
Biological Validation of MPRA Using the *FIRRE* locus. Similar to the MALAT1 Region M and Region E we used to ensure our MPRA was working robustly, we can also use the RRD region from the FIRRE locus. **A**. The MPRA recapitulates the function of known RNA nuclear retention element – RRD. Since, the experiment was performed in human cells, we expect RRD derived from human *FIRRE* to positively influence nuclear enrichment while the RRD derived from mouse *FIRRE* will not influence nuclear enrichment of fsSox2. Here, we show a CDF plot of the nucleotides overlapping human RRD, mouse RRD and other nucleotides in the human and mouse *FIRRE* loci. *P*-value: Mann Whitney Test. **B**. Differential Region-calling correctly identifies nuclear retention elements in *FIRRE*. Solid lines: per-nucleotide abundances in the nuclear (red) and whole-cell (gray) fractions, modeled for each position along the *FIRRE* transcript, based on the aggregate behavior of all oligos containing that nucleotide (*Methods*). Shaded regions: standard deviations. Median values for six biological replicates are shown.

**Figure 6.**
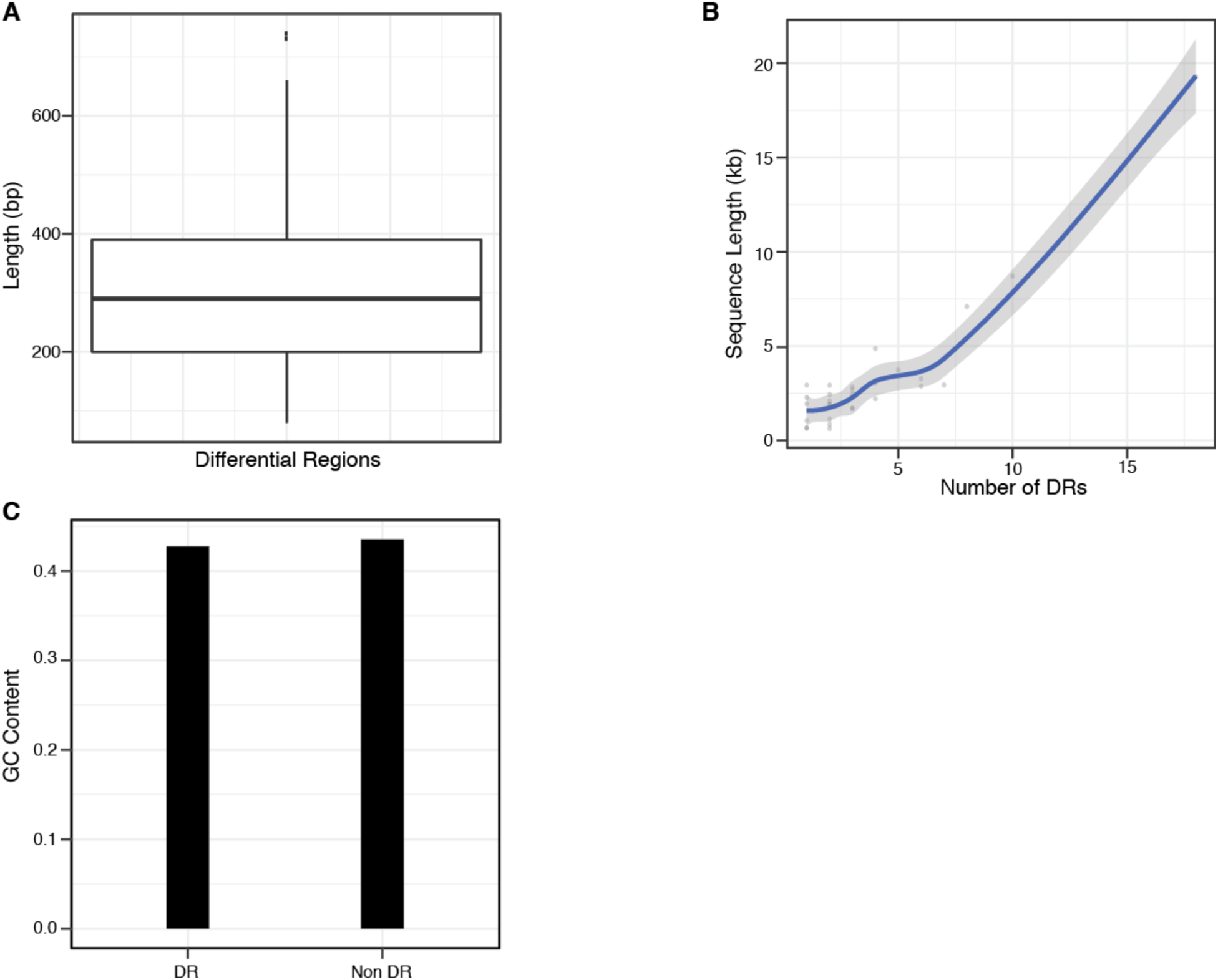
Sequence Features of Differential Regions. **A**. A boxplot showing the length distribution of the differential regions generated by our method. We see that most of our differential regions are longer than 110 bp oligo nucleotide we started with. **B**. A scatter plot showing the relationship between number of differential regions in a lncRNA (X-axis) and the length of the lncRNA (Y-axis). The blue line shows the loess fit and the shaded region is the confidence interval around the fit. **C**. A bar graph comparing GC content of DRs and non DRs which shows there is no noticeable difference in GC content.

**Figure 7.**
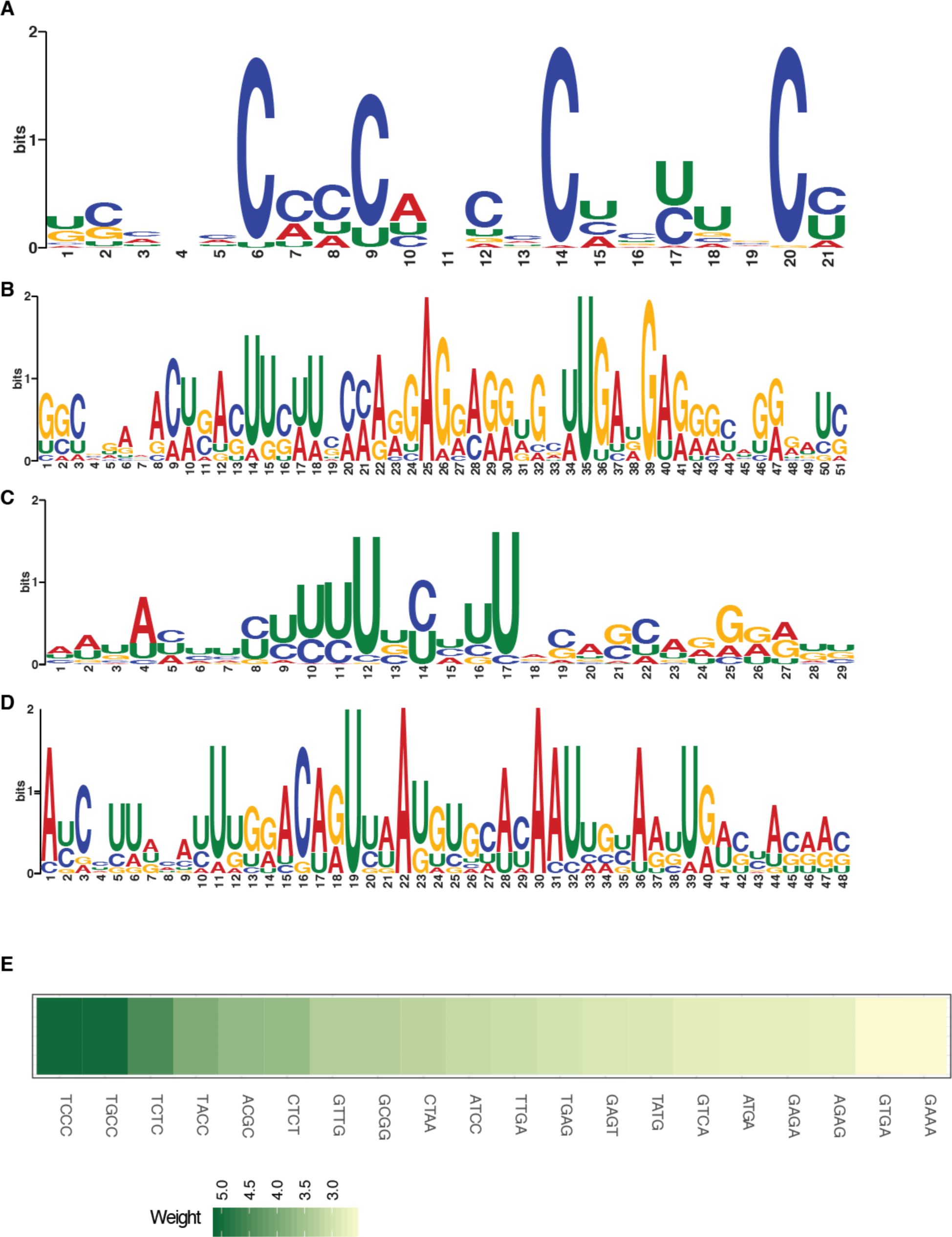
Motifs enriched in lncRNA nuclear enrichment signals. **A-D.** Position Weight Matrix (PWM) for a novel motifs enriched in DR sequences found using MEME software. While motif in panel A is similar to the C-rich motif in Figure 4D the other 3 motifs are found in XIST and similar to the XIST specific motif in Figure 4A E-Value < 0.05. **E**. k-mers mildply predictive of DR found using ridge regression. The color describes the weight of the kmer assigned by the ridge regression algorithm (*Methods*).

**Figure 8.**
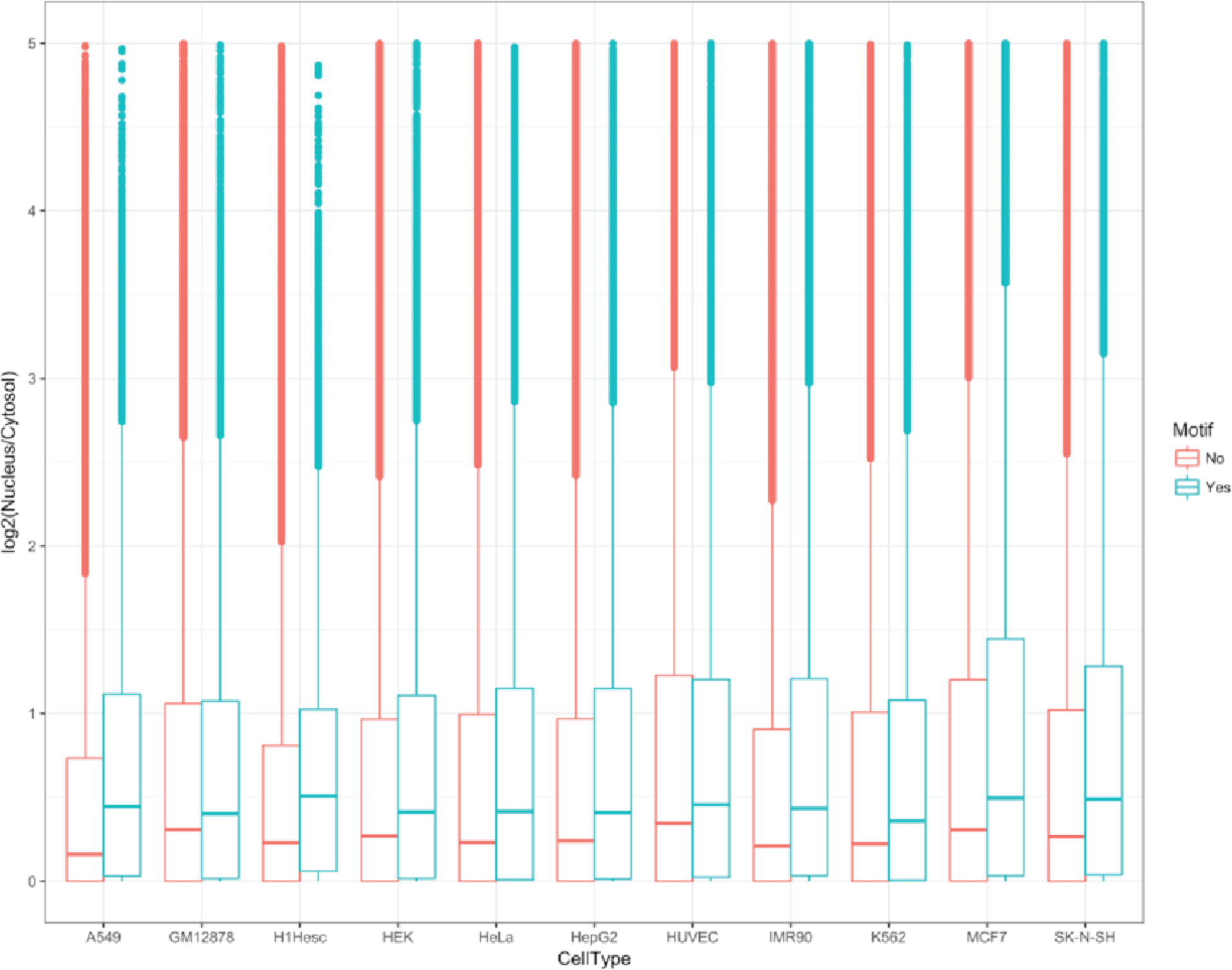
Novel C-rich motif can influence the localization of endogenous human transcripts. CDF plot comparing the nuclear enrichment of all human transcripts with at least one occurrence of our discovered motifs, relative to all other transcripts, in all ENCODE Tier 2 cells^30^. *P*-value: Mann Whitney Test.

